# Identification and Development of Functional Markers Associated with Triple Pistil Phenotype in Wheat

**DOI:** 10.1101/2025.11.09.687415

**Authors:** Qasim Raza, Qilu Song, Shams ur Rehman, Zulfiqar Ali

**Affiliations:** College of Life Sciences, University of Chinese Academy of Sciences, Beijing, China; State Key Laboratory of Wheat Improvement, Peking University Institute of Advanced Agricultural Sciences, Weifang-261325, Shandong, China; Department of Plant Breeding and Genetics, Faculty of Agriculture, University of Agriculture Faisalabad, Faisalabad-38000, Pakistan

**Keywords:** Carpel, CRISPR-Cas9, Floral biology, Grain yield, Marker-assisted selection, Multi-ovary

## Abstract

Triple pistil (TP) wheat is a historical genetic resource capable of producing up to three grains in a single floret and bolstering the current stagnant grain yield potential. TP phenotype is speculated to be the result of a spontaneous mutation; however, the exact underling genetic mechanism remains elusive, with lack of functional markers for early generation trait selection in hybrid wheat breeding programs. Here, scanning electron microscopy highlighted clear developmental differences between single and triple pistil plants started to arise during 1-2 cm long young spike stages. Using a forward genetics approach, we identified consistent TP-associated mutations in two genes (*TraesCS2D02G490900* and *TraesCS2D02G491600*) exhibiting a nearly complete co-segregation with TP phenotype and developed functional markers for early generation trait selection. CRISPR-Cas9 mediated gene-editing of *TraesCS2D02G490900* shifted grain set toward single-grain florets in one edited line in transgenic wheat plants. Furthermore, grain yield evaluation exhibited a significant increase in grains per spike, with no statistically significant reduction in grain weight per spike. Hybrids between common and TP wheat exhibited relatively higher yields, highlighting TP wheat as a significant donor to fortify grain yield potential. This study provides co-dominant functional markers for early generation TP trait selection and valuable targets for hybrid wheat breeding programs.

## Introduction

Bread wheat (*Triticum aestivum* L.) is one of the world’s oldest, most produced, stored, and consumed food crops. Wheat grains comprise a major component of the human diet, supplying nearly 20% of the daily calories and 21% of the protein requirements (Igrejas and Branlard 2020). By 2050, the global human population is likely to surpass 9.7 billion. However, the current rate of genetic gain in wheat production is insufficient to meet the increasing demand driven by global human population growth. The current annual genetic gain in wheat production is estimated to be around 1%, whereas demand is projected to increase by 1.7% (Tadesse et al. 2019; Erenstein et al. 2022). Declining basic resources for crop production, scarcity of fertile lands and alarming increase in population growth rate amounts to a perfect hunger storm. Although, recent advancements in agricultural breeding and technology could potentially accelerate genetic gains beyond current rates, however the challenge remains to implement innovations effectively to meet the rising demands of a growing population.

Wheat yield is a complex trait influenced by multiple agronomic, genetic, environmental, management and their interaction parameters. The genetic yield potential represents maximum yield that a crop can attain under ideal growth conditions e.g., agronomic, environmental and management practices. Grain yield is determined by the number of fertile tillers per plant, the number of spikelets per spike, the number of grains produced per spikelet, and the mean grain weight (Sreenivasulu and Schnurbusch 2012; Sakuma and Schnurbusch 2020). One way to improve grain yield potential is to increase the number of grains per spike, without significant reduction in average grain weight per spike. Floral organs directly influence the number of grains produced and their basic understanding is critical for enhancing the genetic yield potential.

Floral organs are essential for flower architecture, reproduction and grain production, thus providing a basis for grain yield improvement. Wheat spikes are composed of spikelets, and each spikelet bears several florets. Spikelet is the basic unit of a cereal flower (Ali et al. 2019). Each floret contains a lemma, a palea, two lodicules, three stamens and one pistil (alternatives carpel/ovary). Both stamens and pistils are the most critical floral organs and their quantity within each single floret contributes to the final grain yield. An increase in stamens would lead towards the availability of more pollen for efficient pollination and subsequent seed setting. Likewise, an increase in number of pistils/carpels/ovaries would result in higher number of grains per spike, ultimately contributing to the total grain yield potential. Thus, exploration and exploitation of floral modifications related wheat genetic resources are crucial for bolstering the genetic yield potential.

The multi-pistil (alternatives multi-ovary and multi-grain spikelet) phenomenon is common in cereals (Selva et al. 2023; Zhang et al. 2024; Zhang et al. 2025), demonstrating genetic potential to enhance grain yield through production of larger number of grains (Wang et al. 2022). A typical wheat floret produces only a single grain. However, the triple-pistil (TP) wheat germplasm, first reported in 1973 (Chen et al. 1983), contains three pistils or ovaries and produces up to three mature grains within a single floret (Peng 2003). Although, one or two pistils may die out in some florets before seed set due to competition for nutrients, pollen and space, but many florets can set three grains (Raza et al. 2019). TP wheat has normal inflorescence morphology and developmental process just like common hexaploid wheat, except for the occurrence of two extra pistils. The extra pistils are completely fertile, holding ability to produce normal grains and the trait is highly heritable to the next generations (Peng 2003). Until now, several genes regulating the development of floral organs have been reported and functionally characterized in wheat (Ali et al. 2019; Koppolu et al. 2022).

Since publication of first report on TP wheat (Chen et al. 1983), several studies involving TP genotypes originating from genetically different parental materials have been conducted (Peng et al. 2008; Wang et al. 2009; Yang et al. 2017; Zhu et al. 2019; Mahlandt et al. 2021). All of these reports have mapped TP phenotype linked genetic locus on long arm of chromosome 2 of D-subgenome (2DL). Previously, Mahlandt et al., (2021) fine mapped the TP locus (*Mov-1*) on 2DL into a physical region ∼1.1 Mb (589.3 Mbp – 590.4 Mbp) harboring only 26 high confidence genes. More recently, Schoen et al., (2025) reported a map-based identification of *WUSCHEL-D1* (*WUS-D1*) within the *Mov-1* locus as the strong candidate governing multi-ovary phenotype, yet the precise *WUS-D1* knockout in multi-ovary background is still required for validation of causality. Furthermore, the exact underling molecular mechanism remains elusive, with lack of availability of TP or multi-ovary phenotype linked functional markers for early generation selection of trait in hybrid wheat breeding programs.

In this study, we comprehensively investigated TP mutants in spring and winter wheat backgrounds and identified exact stage at which clear developmental differences start to appear between triple pistil and single pistil wheats. Then we used a forward genetics approach to identify potentially mutated genes within *Mov-1* locus and developed candidate functional markers strongly co-segregating with the TP phenotype. We also functionally tested a candidate gene employing CRISPR-Cas9 gene editing. Finally, we made comparisons of grain numbers and grain weight per spike between triple pistil and single pistil recombinant inbred plants to illustrate TP wheat genetic potential. This study advances our understanding on the genetic basis of triple pistil wheat and provides valuable co-dominant functional markers for early-generation trait selection in hybrid wheat breeding programs.

## Materials and methods

### Plant materials

The triple pistil (TP) wheat material was available in spring (STP) and winter (WTP) backgrounds. These lines share similar phenotypic and agronomic traits, except for the spring and winter growth habits. STP seeds were received from CIMMYT, later evaluated, multiplied and collected from the Department of Plant Breeding and Genetics, University of Agriculture Faisalabad (UAF), Pakistan. WTP, previously designated as IGDB-TW (Wang et al. 2009) seeds were collected from the Institute of Genetics and Developmental Biology, Chinese Academy of Sciences, Beijing, China. Chinese spring (CS) and Fielder wheat seeds were collected from the Peking University-Institute of Advanced Agricultural Sciences (PKU-IAAS), Weifang, China. Pirsabak-21 (hereinafter referred as Pirsabak), an elite spring wheat cultivar approved for general cultivation in Pakistan, seeds were also collected from the UAF, Pakistan. Homozygous plants derived from single spikes of each genotype were used for genetic, phenotypic and molecular analyses.

### Growth conditions

Healthy seeds of all genotypes were soaked in 1% hydrogen peroxide (H_2_O_2_) solution for 12 hours, rinsed with tap water for 5 minutes and sown in soil + peat moss mixture containing germinating media in 50 well plastic trays. Ten days old seedlings of spring wheat lines (CS, Fielder, Pirsabak and STP mutant) were directly transferred to flowering pots, whereas winter TP mutant (WTP) seedlings were transferred to 4 °C for vernalization treatment. Vernalized WTP seedlings transferred to flowering pots after 4-5 weeks. All genotypes were grown under controlled growth chamber conditions (22±3 °C, 16 h/8 h light dark cycle, 65±5% relative humidity) with standard management practices, including watering, fertilizing, spraying and weeding. Phenotypic data were recorded at appropriate growth stages. All experiments were performed at the research laboratories and experimental area of PKU-IAAS, China.

### Populations development

Spring TP was reciprocally crossed with Pirsabak, Fielder and WTP. F_1_ plants were confirmed for their hybridity by observing TP phenotype at early anthesis stage and a gene specific marker identified to be polymorphic between parental genotypes. Only confirmed F_1_ plants were allowed to self-pollinate and produce F_2_ segregating populations. Growing of F_2_ plants and their phenotyping were done under controlled environment to minimize environmental variability. Spike phenotypes were precisely recorded from individual F_2_ plants before maturity. Phenotypes of all florets in a main tiller spike were visually recorded. All plants were divided into two distinct groups. Those F_2_ plants which contain at least two florets with three grains were considered as triple pistil (TP), whereas those plants in which all florets of a spike bear single grains were considered as single pistil (SP) plants. This permissive threshold was chosen to minimize false negatives for early-generation selection while recognizing that expressivity varies across florets and spikes.

### Amplification and sequencing of genes

Genomic DNA was extracted from young leaves of healthy plants according to an optimized CTAB protocol. DNA concentration was quantified using a Nanodrop to ensure its high purity (A260/A280 ratio ≥ 1.80). Gene specific primers of 26 high-confidence genes located within *Mov-1* locus (Mahlandt et al. 2021) were designed through PrimerServer online tool at WheatOmics v1.0 database (Ma et al. 2021) and validated at the NCBI Primer-BLAST tool (https://www.ncbi.nlm.nih.gov/tools/primer-blast/index.cgi). PCR reactions were set up with 2x Phanta Max Master Mix Dye Plus (Vazyme, China). PCR products were run on 1.5% agarose gel stained with ethidium bromide to verify amplification and size of expected bands. The amplicons were Sanger sequenced and compared against IWGSC CS Refseq v1.1 (Appels et al. 2018) using SnapGene® (v6.0.2) to identify mutations.

### Marker-trait co-segregation

All F_1_ plants derived from different crosses and a subset of 102 F_2_ individuals derived from STP x Pirsabak cross were genotyped with *TraesCS2D02G490900* and *TraesCS2D02G491600* specific markers. *TraesCS2D02G490900* amplicons from all 102 F_2_ individuals were Sanger sequenced, compared against IWGSC CS Refseq v1.1 and SNPs presence/absence was recorded. Genotypic and phenotypic data were correlated for each F_2_ individual to determine marker-trait co-segregation. The similar approach was adopted for *TraesCS2D02G491600* marker-trait co-segregation analysis.

### Scanning electron microscopy

To collect young spikes of STP and Fielder, the upper portions of representative plants were cut at different developmental stages (Waddington developmental scores 3.0 – 6.0, approx. 30 – 60 days after emergence) (Waddington et al. 1983) and placed in a water filled glass beaker. The stem and leaf tissues enveloping the young spikes were meticulously removed. The freshly harvested young spikes were trimmed to a manageable size (∼5 mm) and immediately immersed in 2.5% glutaraldehyde (in 0.1 M phosphate buffer solution PBS, pH 7.2, Coolaber®, China) for overnight at 4 °C. After overnight fixation, samples were rinsed 3 times (10 minutes each) with 0.1 M PBS to remove excess fixative. Then samples were dehydrated using an ethanol series: 30% for 10 minutes, 50% for 10 minutes, 70% for 10 minutes, 90% for 10 minutes and 100% for 10 minutes (2 times). After complete dehydration, samples were transferred to a critical point dryer using 100% ethanol. The dried sample were mounted on SEM stubs using carbon tape and gold coated using a sputter coater. Finally, coated samples were loaded into the SEM chamber. The developmental morphology of young florets and floral primordia at different stages was observed and documented via Helios 5 CX DualBeam scanning electron microscope (ThermoFisher Scientific).

### CRISPR-mediated knockout of candidate gene

Single guide RNA (sgRNA) target site in exon of candidate gene was retrieved from CRISPR-Cereal (http://crispr.hzau.edu.cn/cgi-bin/CRISPR-Cereal/main) online web tool. The gene specific forward and reverse sgRNA target sequences were synthesized and annealed to form double-stranded oligonucleotide adapters. sgRNA was PCR connected to the wheat TaU3 promoter. TaU3-sgRNA cassette was cloned into a CRISPR/Cas9 vector backbone (CasV2-GRF4-GIF1) through InFusion cloning. The cloning process was completed using ClonExpress II One Step Cloning Kit (Vazyme, China) by following manufacturer protocol. The cloned product was introduced into Trelief® 5α chemically competent cells (*E. coli*) and individual positive colonies were identified through colony PCR. The intermediate recombinant plasmids were purified using FastPure EndoFree Plasmid Mini Plus Kit-Box2 (Vazyme, China) and introduced into the EHA105 chemically competent cells (*Agrobacterium tumefaciens*) following manufacturer protocols. Finally recombinant CRISPR-Cas9 plasmid was introduced into the STP mutant through *Agrobacterium*-mediated transformation (Ishida et al. 2015). The CRISPR-Cas9 edited lines were generated by following the gene editing technology in wheat (Debernardi et al. 2020). Regenerated T_0_ generation plants were screened for the presence of Cas9 carrier. DNA extraction and amplification of target gene regions were completed using gene specific primers. PCR products were analyzed by gel electrophoresis and Sanger sequenced to confirm presence of indels introduced by CRISPR-Cas9. Homozygous gene edited plants were subsequently identified in the T_1_ generation. The T-DNA/Cas9 carrier free plants were identified in the T_1_ generation using transgenic Pat/Bar speed test strips (Wuhan Apbios, China).

### Statistical analysis

All experiments were conducted in triplicate at least. GraphPad Prism (v.10.2.3) was used for data analysis. Phenotypic data were compared following an ordinary ANOVA and Tukey’s multiple comparisons test. Significance among groups was determined based on a non-parametric Kruskal-Walli’s test.

## Results

### Phenotypic characterization of TP mutants and SP wheat genotypes

The widely common single pistil (SP) cultivars and the less common triple pistil (TP) mutant genotypes were grown under controlled growth chamber conditions (22±3 °C, 16 h/8 h light dark cycle, 65±5% relative humidity) (**Fig. 1a**). The TP mutants (STP and WTP) and SP genotypes (Fielder and Pirsabak) share nearly similar inflorescence morphology and developmental processes, except for the occurrence of one or two additional pistils/ovaries in developing florets of TP mutants (**Fig. 1c-f**). Although, one or two pistils may fail to develop in some florets due to competition for nutrients, pollen and space (**Fig. 1e-f**) , however majority of the florets contain double or triple pistils. The additional pistils are completely fertile holding ability to produce normal grains in back-to-back fashion (**Fig. 1g**). Additionally, TP spikes are longer and wider than SP spikes, along with sterility of terminal spikelets (**Fig. 1a**). Apart from the distinct floret morphology, in our materials TP mutants are taller, with longer peduncles and produced relatively fewer tillers than SP genotypes (**Fig. 1h-k**). Furthermore, TP mutants exhibit stickiness of their spikes, attract more insects and pests, and flower earlier than SP genotypes. These distinct morphometric characteristics highlight TP wheat as a valuable genetic resource for wheat genetics and breeding.

**Fig. 1.**
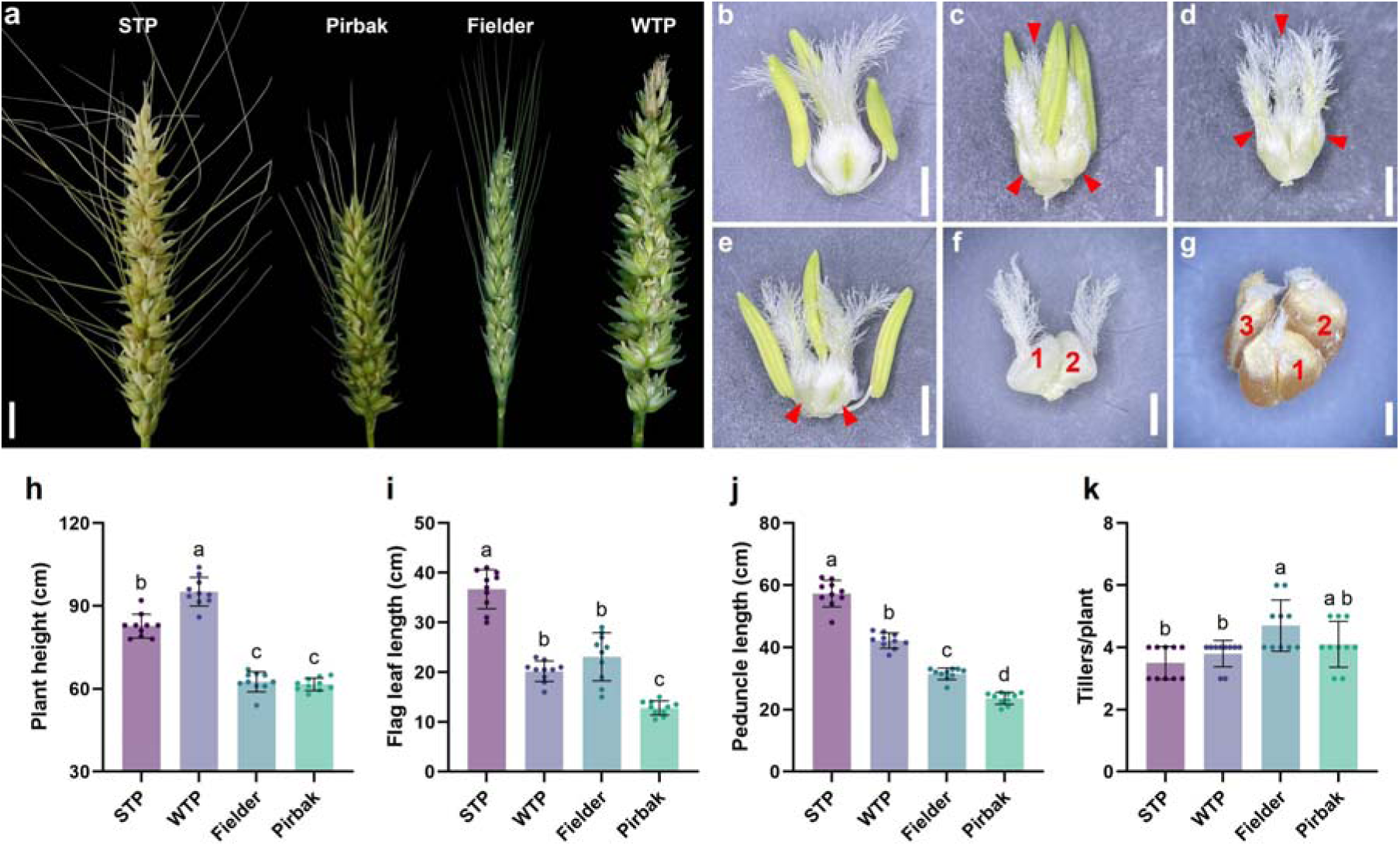
Phenotypic characterization of triple pistil and single pistil wheat genotypes. (**a**) spikes at grain formation stage. Scale bar, 1 cm. (**b**) Fielder and (**c-f**) STP reproductive floral organs (stamens and pistils) within single florets. Scale bars, 1 mm. (**g**) Three close set grains in STP floret. Scale bar, 5 mm. (**h-k**) Plant height, flag leaf length, peduncle length and number of tillers per plant comparison among TP and SP genotypes. Multiple comparisons were determined based on an ordinary ANOVA and Tukey’s test.

### Scanning electron microscopy of developing young spikes

To identify the exact developmental stage at which noticeable phenotypic differences between TP and SP floral organs begin to appear, the Fielder and STP young spikes were examined under scanning electron microscope at different developmental stages. In both genotypes, the floret meristem primordia start to differentiate into floral organs at stage 1, followed by the complete distinction of a lemma, a palea, three stamens and a pistil primordium at stage 2 (**Fig. 2**). However, additional pistil primordium starts to emerge from the base of main pistil and adjacent to the lemma in the STP florets at stage 3, whereas Fielder florets lack an additional pistil primordium and possess only a single main pistil primordium at this stage. In both genotypes, the reproductive floral organs continue to fully differentiate into the three stamens and one main pistil primordium from stage 3 onward. However, only in STP did the additional pistil primordia continue differentiation during stages 3 and 4, lagging behind the main pistil but ultimately developing into the triple pistil floret (**Fig. 2**). This developmental mechanism exposed that clear floral differences between TP and SP florets start to appear when the developing young spikes are 1 – 2 cm long and this developmental stage is good for subsequent experiments.

**Fig. 2.**
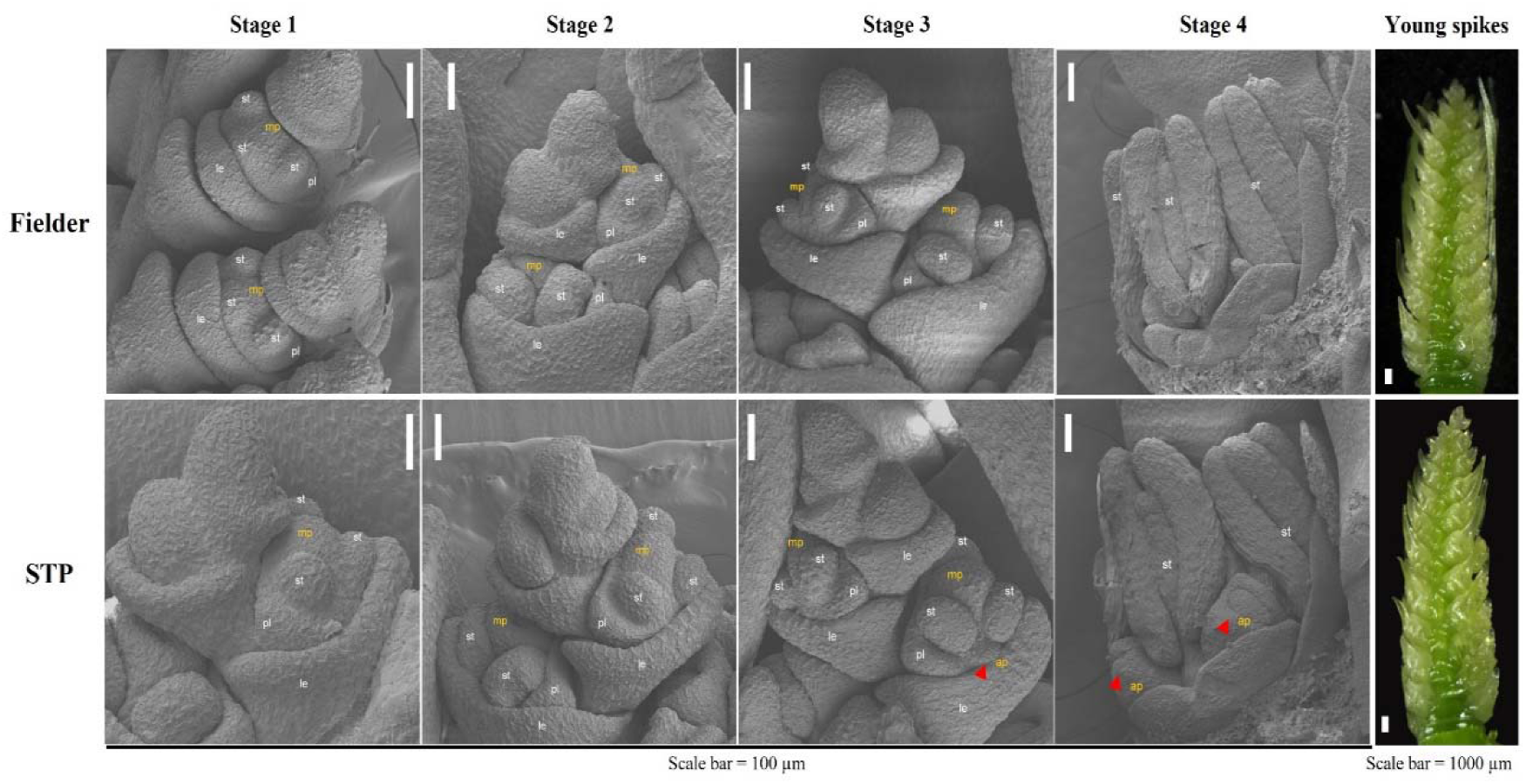
Comparative scanning electron microscopy of young spikes at different developmental stages. Stages 1 – 4 indicate the four stages of floret developmental processes within 1 – 2 cm young spikes. For both genotypes, 100 spikelets from 10 individual plants were observed. The red triangle represents the additional pistils that develop into the mature ovaries and grains. le, lemma; pl, palea; st, stamen; mp, main pistil; ap, additional pistil.

### Inheritance of TP trait

To investigate genetic basis of the TP trait, STP mutant was crossed with two SP cultivars (Fielder and Pirsabak) to develop independent segregation populations (**Fig. 3** & **S1a-b**). Additionally, STP and WTP mutants were crossed with each other to assess potential allelism for TP trait. All F_1_ plants exhibited the TP trait with similar percentages to their TP parental genotypes, except for the F_1_ derived from STP x WTP cross, which exhibited a significantly higher percentage of TP trait (**Fig. 3a**). In the two reciprocal F_2_ populations derived from STP x Fielder and STP x Pirsabak crosses, the mendelian segregation ratio of TP to SP plants conformed to a 3:1 ratio (χ²0.05 = 1.48, 0.91) (**Fig. 3b**). In the STP x WTP derived F_2_ population, all plants exhibited the TP trait. These results are consistent with a single dominant nuclear locus and the distinctive phenotype of STP and WTP plants is determined by the same locus.

**Fig. 3.**
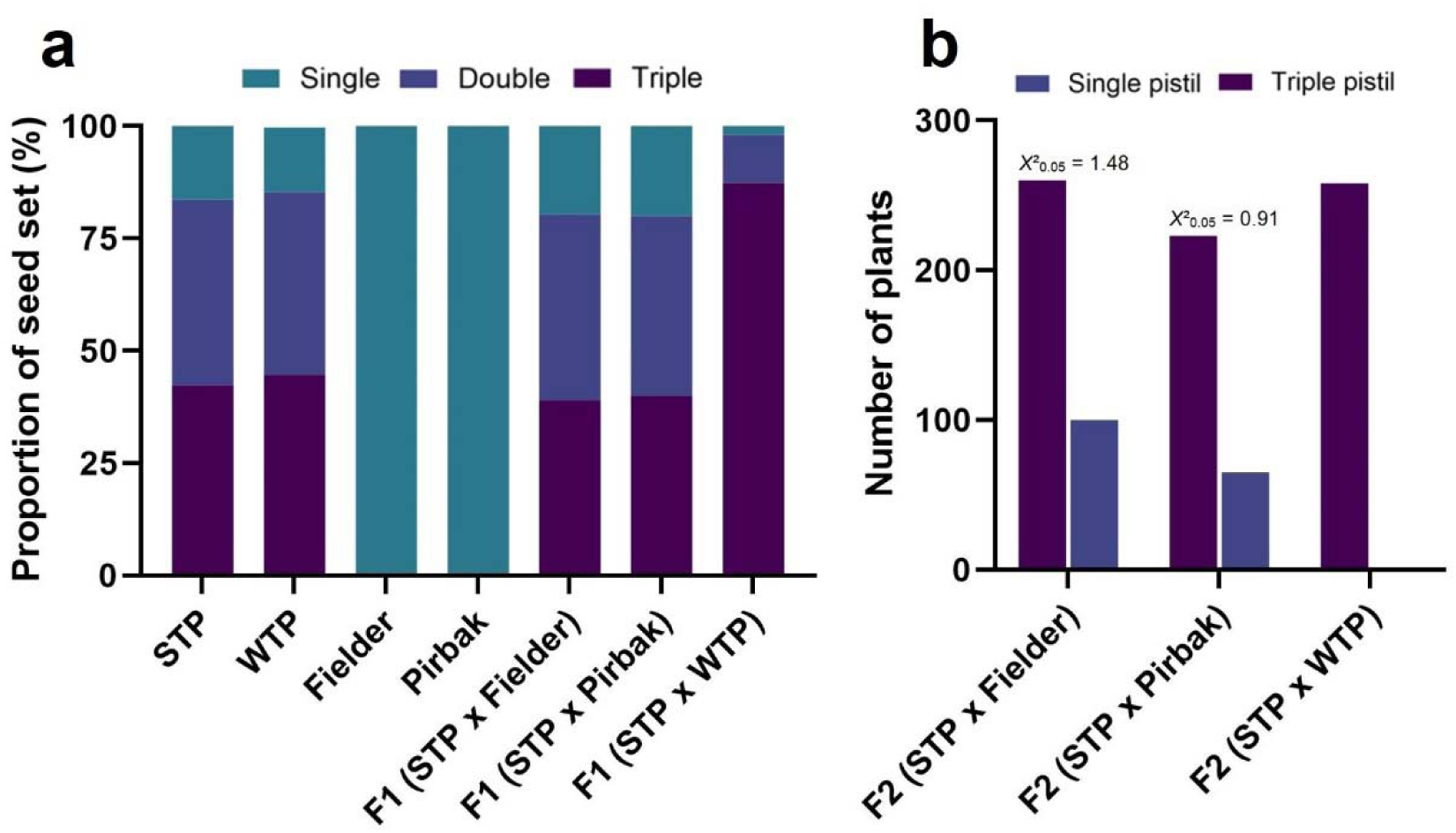
Genetic analysis of triple pistil trait. (**a**) Proportion of single, double and triple grain florets in parental and F1 genotypes. Mean data derived from three independent main tiller spikes. (**b**) Phenotypic segregation in F_2_ populations.

### Identification of mutated genes within Mov-1 locus

*Mov-1* locus was previously fine mapped to a physical interval of ∼1.1 Mb (589.3 Mb – 590.4 Mb) on distal end of 2DL chromosome (Mahlandt et al. 2021). This physical interval harbors only 26 high confidence protein encoding genes (**Fig. 4**). Amplicon-based screening of high-confidence genes within fine-mapped intervals is a cost-effective first step to deliver functional markers for breeding. Taking advantage of this mapping data, gene specific primers were directed to 2D and validated using Primer-BLAST. The amplicons were Sanger sequenced and compared against IWGSC CS Refseq v1.1. In our amplicon survey, two genes (*TraesCS2D02G490900* and *TraesCS2D02G491600*) showed consistent TP-associated variations (**Fig. 4a**). *TraesCS2D02G490900* had 21 SNPs in protein coding sequence (corresponding to 9 non-synonymous amino acid alterations) of STP and WTP mutants and encodes an agmatine coumaroyltransferase protein (**Fig. 4b**). Its orthologs in *Arabidopsis* regulate cell expansion and plant growth via phenolic precursors and auxin transport (Abdulrazzak et al. 2006; Besseau et al. 2007). *TraesCS2D02G491600* encodes a leucin-rich repeat receptor like protein kinase (LRR-RLK), showed amplicon loss in coding and UTR regions, consistent with a deletion in TP materials (**Fig. 4c**). RLK is a large family of proteins with over 929 members in hexaploid wheat and several of these proteins are reported to regulate inflorescence development and immunity (Cui et al. 2022; Liu et al. 2023; Yu et al. 2025).

**Fig. 4.**
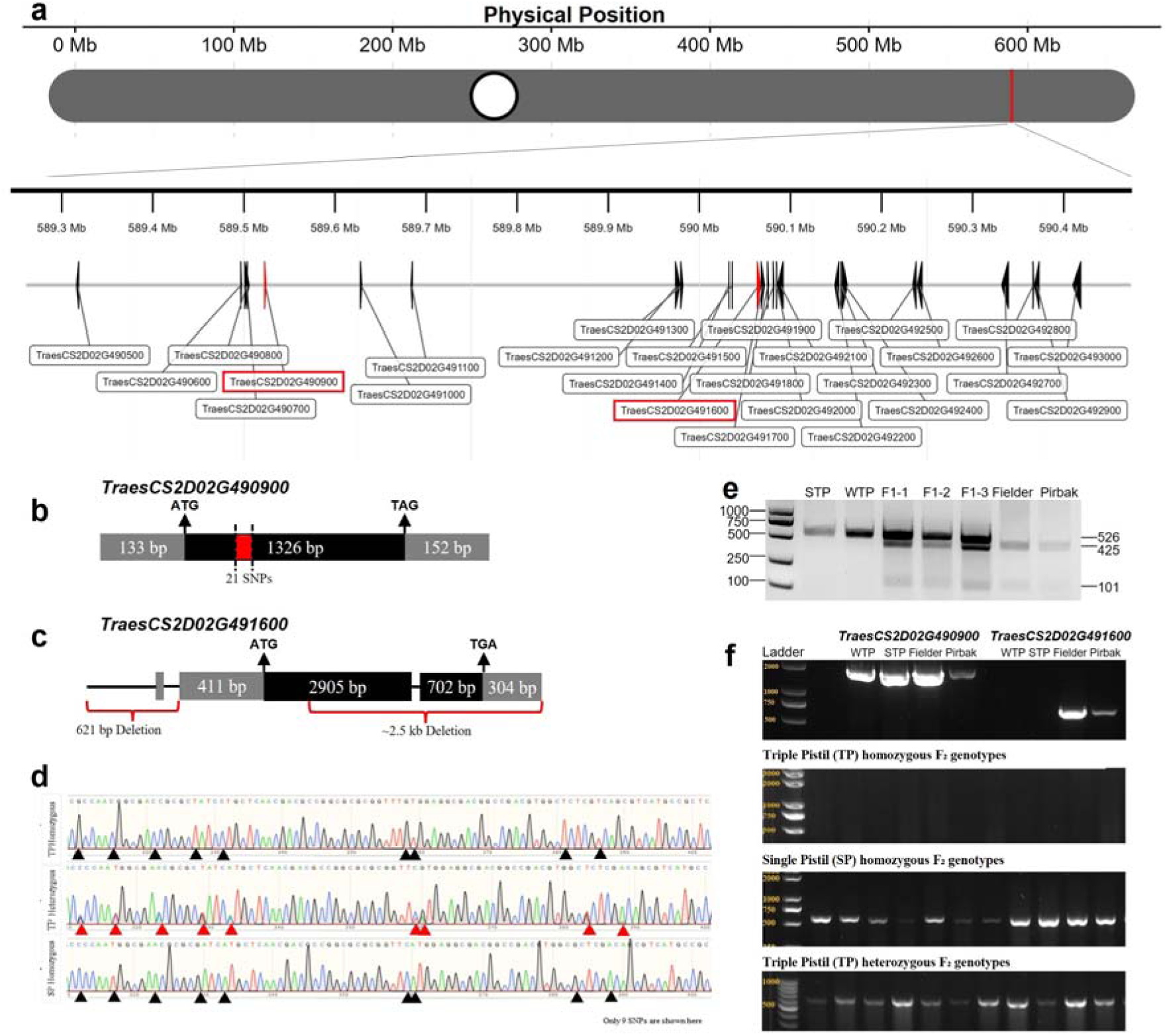
Chromosomal location of *Mov-1* locus and TP phenotype linked genes. (**a**) Physical position of *Mov-1* locus on 2DL, encompassing 26 high confidence genes, (**b**, **c**) schematic diagrams of mutated genes. Exon and intron are indicated with black box and solid line, respectively. 5’ and 3’ untranslated regions (UTRs) are indicated with gray boxes. SNPs and deletions are represented with red box and brace symbols, respectively. (**d**) Polymorphic SNPs among parental and heterozygous F_2_ genotypes. Black triangles in TP and SP homozygous genotypes indicate SNPs between parental STP and Pirsabak plants. Red triangles in heterozygotes indicate the presence of both types of nucleotides from STP and Pirsabak parents. Only 9 SNPs are shown here for simplicity. (**e**) Development of *TraesCS2D02G490900* specific CAPS marker. (**f**) Gel-electrophoresis graphs of *TraesCS2D02G491600* deletion specific functional marker. Upper left and right panels represent gDNA amplification from parental genotypes using a positive control marker (*TraesCS2D02G490900*) and RLK-deletion specific marker (*TraesCS2D02G491600*), respectively. Lower panels represent gDNA amplification from TP homozygous, SP homozygous and TP heterozygous F_2_ genotypes.

### Validation of Mov-1 locus genetic mapping and development of functional markers

Since *Mov-1* locus was previously reported, we reasoned to validate the genetic mapping by marker-trait co-segregation analysis before proceeding with further analyses. To evaluate this, a cross was made between STP mutant and Pirsabak wheat cultivar (**Fig. S1a**). The resultant F plants were allowed to self-pollinate and produce F population. Before self-pollination, hybridity of F plants was confirmed through phenotypic (TP phenotype before anthesis) and genotypic (presence of 21 SNPs in TP plants) characterization. A subset of 102 F plants were grown under controlled glasshouse conditions by following standard agronomic and plant protection practices. Out of total 102, 71 plants showed TP phenotype, and 31 plants showed SP phenotype, indicating a 3:1 Mendelian segregation (χ²0.05 = 1.9075). PCR-amplification and sanger sequencing of all 102 F plants with *TraesCS2D02G490900* specific primers revealed strong marker-trait co-segregation (> 97%) (**Table 1**, **Fig. 4d**), validating that previous genetic mapping report on *Mov-1* locus is reliable. Additionally, PCR-amplification of parental genotypes and F_2_ plants with a *TraesCS2D02G491600* deletion-specific primer pair also revealed strong marker-trait co-segregation (> 96%) with TP phenotype (**Fig. 4f**). Small variations in marker-trait co-segregation could be explained by environmental variables. These results highlight that previous genetic mapping of *Mov-1* locus is reliable, and the TP trait governing gene or genomic region is positioned within the same locus.

**Table 1.** Marker-trait co-segregation in a subset of F_2_ population (STP x Pirsabak)

| **Description** | **Triple Pistil**<br>**(TP)** | **Single Pistil**<br>**(SP)** | **Co-**<br>**Segregation** |
| --- | --- | --- | --- |
| Phenotype in F<sub>2</sub> | 71 | 31 | > 97% |
| Genotype in F<sub>2</sub><br>(*TraesCS2D02G490900*) | 74<br>20 (Homozygous)<br>54 (Heterozygous) | 28<br>(Homozygous) |  |
| Genotype in F<sub>2</sub><br>(*TraesCS2D02G491600*) | 70<br>20 (Homozygous)<br>50 (Heterozygous) | 28<br>(Homozygous) | > 96% |

Since the two mutated genes exhibited nearly complete co-segregation with TP trait, we reasoned to develop functional markers for these mutated genes. During populations development, spikes of all F_1_ plants derived from different crosses uniformly exhibited TP trait (**Fig. 3a** & **S1a-b**). Genotyping with *TraesCS2D02G490900* and *TraesCS2D02G491600* specific markers confirmed heterozygosity of F_1_’s derived from STP x Fielder and STP x Pirsabak crosses (**Table S1**). Whereas STP x WTP derived F_1_’s showed homozygosity for the two gene-specific markers. For *TraesCS2D02G490900*, we developed a cleaved amplified polymorphic sequence (CAPS) marker by amplifying the coding region encompassing 21 SNPs and enzymatic digestion of different amplicons with *SacI* restriction enzyme (**Fig. 4e**, **Table S1**). Similarly, a deletion-specific marker was developed within coding and 3’ untranslated regions (UTR) of *TraesCS2D02G491600* (**Fig. 4f**). In the STP x Pirsabak F , both the *TraesCS2D02G490900* CAPS and the *TraesCS2D02G491600* deletion assays showed near-complete co-segregation (>96-97%) with the TP phenotype (**Table 1**). Both of these co-dominant functional markers can efficiently differentiate between homozygous and heterozygous TP genotypes during early generation trait selection.

### CRISPR-mediated knock-out of candidate gene

Due to the observed variations at the DNA level along with strong marker-trait co-segregation, *TraesCS2D02G490900* was selected for CRISPR-Cas9-mediated knock-out. Transgenic plants were generated in STP mutant background facilitated by the *Agrobacterium*-mediated genetic transformation (**Fig. 5a**). Two homozygous *TraesCS2D02G490900*-gene edited lines free from the Cas9 carrier were obtained in the T_1_ generation. The first line (T1-3-9) exhibited 7 bp deletion resulting in the acquisition of a premature stop codon at +128 amino acids, as well as frame shift mutation of 49^th^ to 128^th^ amino acids. The second line (T1-9-1) exhibited 15 bp deletion resulting in truncation of 5 amino acids (+46 to +50) along with a substitution of alanine at +51 into tyrosine (**Fig. 5a**). At maturity, the majority (∼79%) of florets in T1-3-9 line displayed both double and triple grains, nearly the same as the non-transgenic STP line (WT). Whereas > 86% of the florets in T1-9-1 line contained single grains, compared with ∼19% in WT line (**Fig. 5b**). However, phenotypic examination of another two homozygous gene-edited lines positive for Cas9-carrier, containing premature stop codons as well as frame shift mutations, indicated a considerable percentage of florets containing both double and triple grains, consistent with the WT line. We thus concluded that although *TraesCS2D02G490900* exhibited a nearly complete co-segregation with TP trait, however, it might not govern the TP phenotype alone and require a non-functional *TraesCS2D02G491600* and/or some other genes. Knock-out of *TraesCS2D02G491600* in WT background (single pistil common wheat) could reveal if this gene governs the TP phenotype or not. Nevertheless, we speculate that the causal gene or genomic region is located within the *Mov-1* locus and has strong linkage with *TraesCS2D02G490900* and *TraesCS2D02G491600*.

**Fig. 5.**
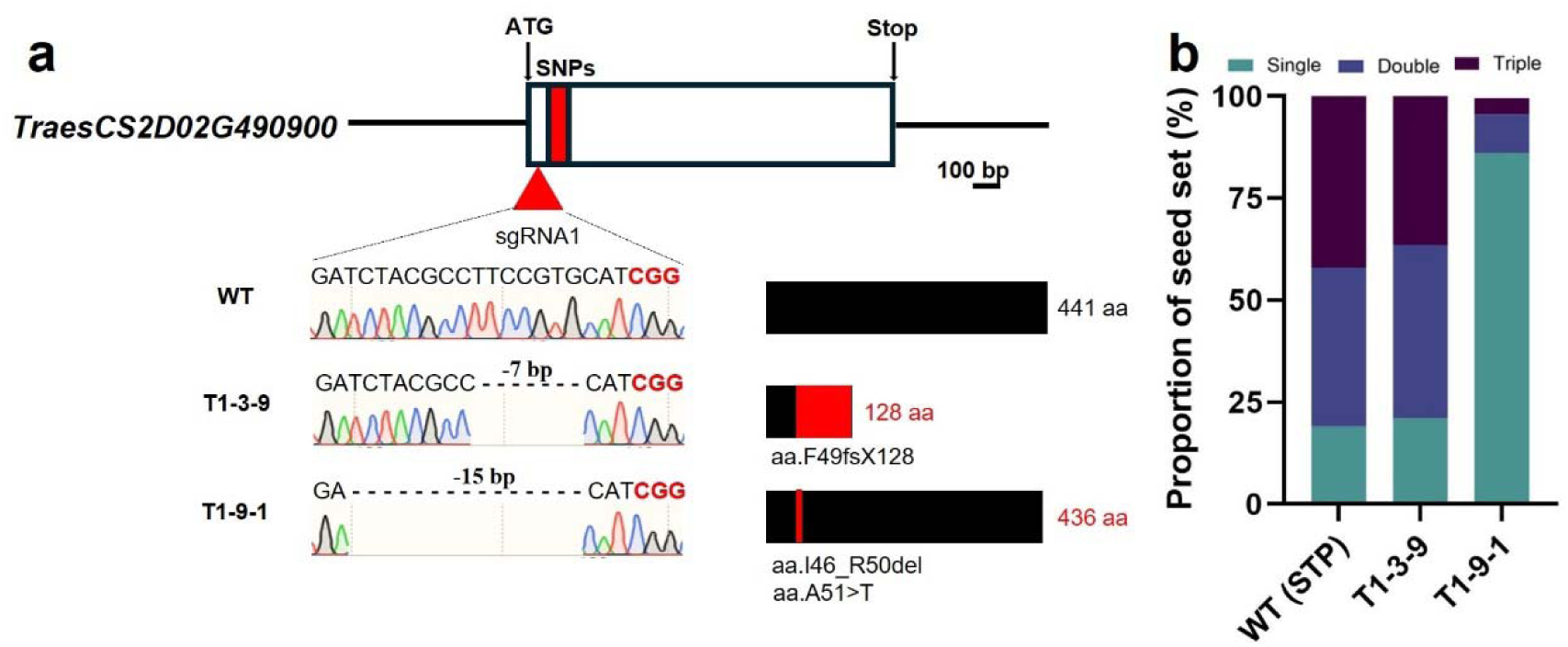
CRISPR-mediated knock-out of *TraesCS2D02G490900*. (**a**) Schematic representation of sgRNA design and CRISPR-Cas9 edited lines along with the resultant proteins. (**b**) Proportion of single, double and triple grains in transgenic and WT lines at maturity stage. Mean seed setting data from two whole spikes including the main spike.

### Grain yield potential of TP trait and prospects for hybrid wheat breeding

To evaluate the grain yield potential of TP trait, we examined the grains per spike and grain weight per spike traits among individuals of an F_2_ population (STP x Pirsabak) (**Fig. 6a**). The results revealed a highly significant (p = 0.0001; p = 0.0005) increase in grains per spike of TP individuals as compared with SP individuals (**Fig. 6b**), and non-significant comparisons for grain weight per spike (p = 0.4885; p = 0.0718) (**Fig. 6c**). It is worth mentioning that several TP individuals exhibited an increase in grain numbers per spike along with grain weight per spike as compared with SP individuals. In general, heterozygous individuals possess genetic potential to increase the grain yield of wheat hybrids. Collectively, these data show increased grains per spike in TP individuals, without a statistically significant reduction in grain weight in the glasshouse F_2_ cohort. However, hybrids derived from TP and SP wheats can produce higher grain yield. These observations illustrate TP wheat as a significant donor to fortify an important yield-related component, along with potential implications in hybrid wheat breeding by increasing grain number per spike and reducing the cost of hybrid seed production.

**Fig. 6.**
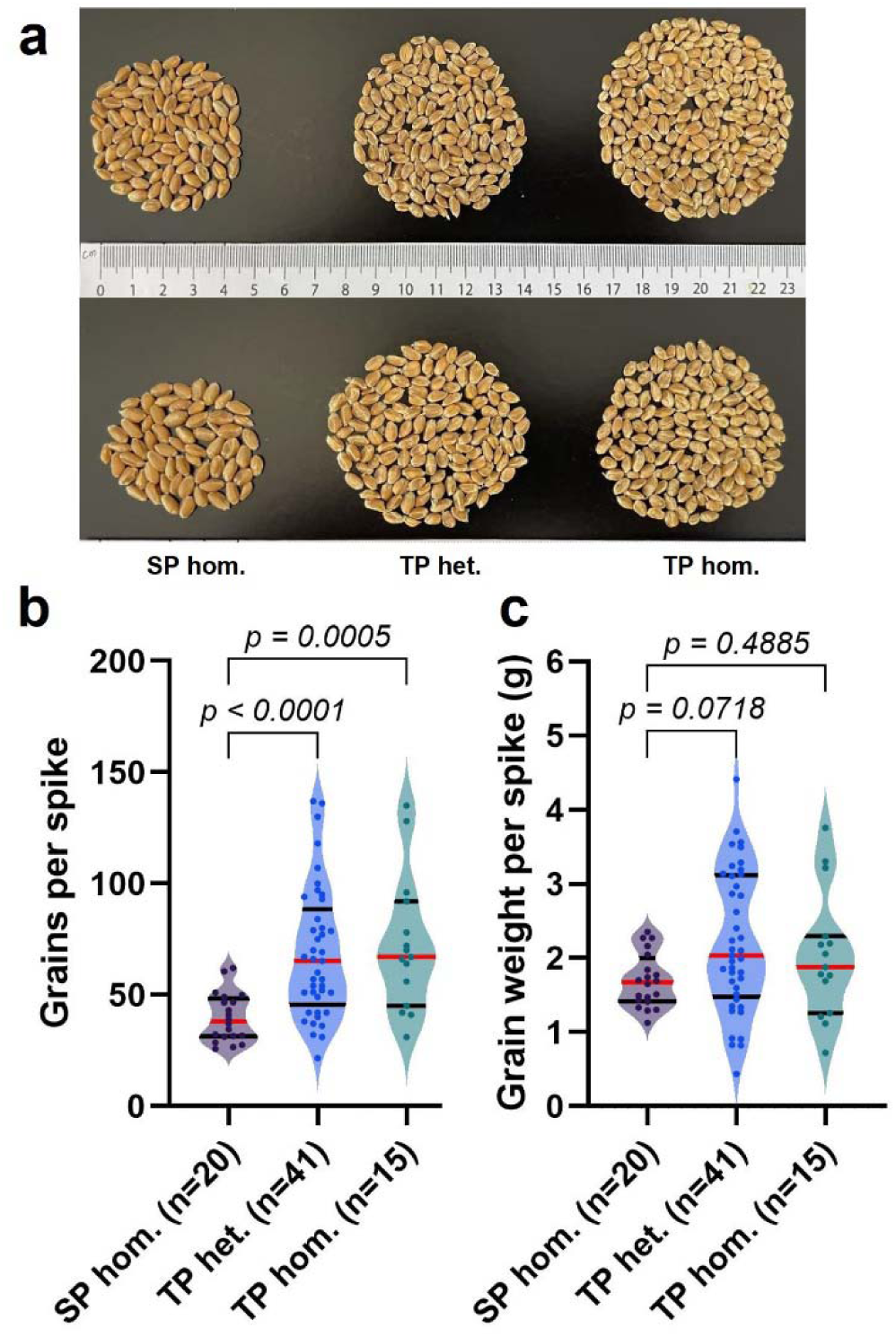
Grain yield potential of TP trait. (**a**) Grains per spike among representative individuals of an F_2_ population derived from STP x Pirsabak cross. Comparison of grains per spike (**b**) and grain weight per spike (**c**) traits among F_2_ individuals. Solid black lines indicate 25% and 75% quartiles, whereas solid red lines indicate median values. Significance among groups was determined based on a non-parametric Kruskal-Wallis test using GraphPad prism.

## Discussion

Bread wheat is critical for global food security. However, its production is stagnant over the past few decades, requiring exploitation of innovative approaches and available genetic resources for further genetic improvements (Schauberger et al. 2018). Grain yield is a complex trait affected by several genetic, environmental and management factors. Among these factors, the source-sink relationship directly influences the crop yield potential. Source is a material producer and exporter e.g., roots, leaves and stems to some extent; while sink is a material importer and consumer e.g., spike, spikelets and grains. A well-coordinated source-sink relationship could realize the promised increase in crop yield potential (Chang and Zhu 2017). Several approaches have been proposed to enhance source-sink capacity and subsequent yield potential. Among these, an improved sink capacity (number of grains produced per unit of spike, grain size and weight) is frequently integrated as a selection target in wheat breeding programs (Hu et al. 2016; Simmonds et al. 2016; Adamski et al. 2021). Although research efforts have yielded significant improvements in wheat sink capacity (Zhang et al. 2022; Zhou et al. 2023), however, progress on increasing the number of grains per unit spike, especially the per individual spikelet, is limited. Furthermore, mystery behind the genetic basis of triple pistil phenotype remains obscure, with non-availability of trait linked functional markers for early generation selection in wheat breeding programs.

In the present study, we investigated triple pistil (TP) and single pistil (SP) wheat genotypes and found that TP wheat has relatively better source-sink capacity (longer flag leaves and stems, increased number of grains and bigger spikes) as compared with SP wheat (**Fig. 1**), which could be further improved for a well-coordinated source-sink interaction and subsequent plant architecture as well as grain yield potential. Since, in the current study architectural comparisons were made between distinct genotypes, we cannot ascribe height, peduncle length and tiller number differences solely to the TP locus without within-family controls. Scanning electron microscopy of developing young spikes highlighted distinct developmental perturbations among TP and SP florets during 1-2 cm long spike stages, where additional pistil primordia start to originate from the base of main pistil and develop into mature ovaries and grains upon plant maturity (**Fig. 2**). TP and SP genotypes derived segregating populations indicated that TP trait is controlled by a single dominant locus, with spring triple pistil (STP) and winter triple pistil (WTP) genotypes conformed to be allelic to each other (**Fig. 3**). These results are consistent with previous reports (Peng 2003; Wang et al. 2009; Mahlandt et al. 2021) confirming that TP trait is governed by a single nuclear locus. However, as the binary TP/SP grouping aggregates heterogeneous expressivity, the 3:1 segregation pertains to the presence of any TP florets under our scoring rule. Future analyses will quantify TP as a portion trait across multiple spikes to refine genetic inference and minimize classification bias.

Previously, multiple studies have mapped TP trait on chromosome arm 2DL (Peng 2003; Peng et al. 2008; Wang et al. 2009; Yang et al. 2017; Zhu et al. 2019; Mahlandt et al. 2021), except one study which placed it on chromosome 5B (Peng et al. 2004). These studies relied on F_2_ segregating populations and employed SSR, GBS and KASP markers, as well as radiation hybrid technique, for genetic mapping of TP locus. Yu et al. (2021) constructed the first partial genetic linkage map using bulked segregant analysis and narrowed down the *Pis1* locus at a genetic distance of 0.6 cM. Later, Mahlandt et al. (2021) fine mapped the *Mov-1* locus within a ∼1.1 Mb physical interval, encompassing 26 high confidence protein encoding genes. In the present study, we conducted amplicon-based screening within *Mov-1* locus and compared amplicon sequences against IWGSC CS RefSeq v1.1 (**Fig. 4**). Our amplicons primarily covered exons and proximal untranslated regions; intergenic regions within *Mov-1* were not comprehensively interrogated. We found consistent TP-associated mutations in two genes (*TraseCS2D02G490900* and *TraesCS2D02G491600*), confirmed the validity of *Mov-1* locus with mutated genes specific markers and phenotyping of F_2_ individuals under controlled glasshouse conditions. Our marker-trait co-segregation analysis revealed near complete co-segregation (**Table 1**), indicating that TP phenotype responsible gene is located within the *Mov-1* locus and have strong linkage with the two mutated genes. Recently, Yamamoto et al. (2023) identified two genes (*TraesCS2D02G491200* and *TraesCS2D02G491600*) which showed downregulation in the three-pistil mutant using a comparative RNA sequencing approach. These two genes are also located within the *Mov-1* locus. Additionally, these researchers found that *TraesCS2D02G491200*/*ARF5* got deleted in the three-pistil mutant background. In this study, we also found that *TraesCS2D02G491600* got deleted in the TP mutant background. It is noteworthy that previous studies used either winter or spring triple pistil mutants. Here, we investigated both mutants at the same time for a more comprehensive analysis. Although we incompletely amplified majority of the genes within *Mov-1* locus, however, only 16 genes were consistently amplified in both STP and WTP mutants, with at least 10 genes being amplified in only one mutant at a time. Based on these observations, we speculate that *Mov-1* locus might contain large structural variation (SV) leading to the deletion of few genes. More recently, Schoen et al., (2025) reported a chromosome-level genome assembly of multiovary wheat that showed large SVs within the *Mov-1* locus, resulting in the deletion of 10 genes, including *TraesCS2D02G491200*/*ARF5* and *TraesCS2D02G491600/RLK*. These results are consistent with findings of current study, indicating complex SVs deriving the formation of multiovary-producing florets. In future, it will be interesting to knockout and phenotype the *WUS-D1* in TP background and large SV region in SP background Click or tap here to enter text.to precisely identify either *WUS-D1* upregulation or large SV is behind the gain-of-function of TP phenotype.

Functional markers derived from polymorphic sites and SVs tightly linked with phenotypic trait variation are considered ideal for marker-assisted breeding (Andersen and Lübberstedt 2003). In wheat, development of gene-derived functional markers is challenging because of polyploid nature (Bagge et al. 2007). To date, none of the TP phenotype linked functional marker is available, hindering robust screening and exploitation of TP trait in wheat molecular breeding programs. We identified and developed CAPS (*TraesCS2D02G490900*) and InDel (*TraesCS2D02G491600*) functional markers strongly associated with the TP phenotype (**Fig. 4e-f**). These co-dominant markers are robust, cost-effective and can be easily implemented in the laboratory. Their potential values for selection of TP trait were validated by marker-trait co-segregation analysis (**Table 1**). These candidate functional markers co-segregate with TP in the STP x Pirsabak cross and can be used for early generation selection of TP trait in wheat breeding programs aimed at introgression of TP phenotype and increasing number of grains per unit spike. However, as our results derive from a single F_2_ background, linkage decay and structural variation around *Mov-1* may differ across germplasm, potentially affecting marker performance. Therefore, broader validation across independent populations and diverse panels is required before routine deployment.

CRISPR-Cas9 has revolutionized plant functional genomics by providing a highly precise, efficient, and versatile toolkit to directly link genes to their biological functions. It is the cornerstone of modern plant biology research. *Arabidopsis* orthologs of the two mutated genes identified in the present study are reported to regulate cell expansion and plant growth (Abdulrazzak et al. 2006; Besseau et al. 2007; Cui et al. 2022). We generated *TraesCS2D02G490900* gene-edited lines with CRISPR-Cas9 to reveal its regulatory roles in TP phenotype (**Fig. 5**). However, floret phenotypes of homozygous knock-out lines exhibited marginal rescue of wild type single pistil phenotype, suggesting other gene(s) might govern TP phenotype. In future, it will be interesting to precisely identify and functionally characterize the causal gene through CRISPR-Cas9, overexpression and complementation experiments.

Floral modifications provide a unique resource for understanding and manipulating yield-contributing factors. Redesigning the wheat flower has potential to facilitate hybrid wheat development and grain yield improvement (Whitford et al. 2013; Selva et al. 2020; Rohde et al. 2025). Identification, cloning and characterization of genes regulating floral development hold enormous potential to engineer higher-yielding wheat cultivars. Previous studies have shown that hybrids between normal and TP/MOV-wheat lines result in substantial yield improvements (Skovmand et al. 2001; Mahlandt et al. 2021)). In this study, we also observed enhanced yield in such hybrids (**Fig. 6**). Although, our results revealed a significant increase in grains per spike along with a non-significant increase in grain weight per spike, however, several hybrids produced higher grain yield. The observed trade-off between grain numbers and weight might be explained by ‘sink-source’ capacity and could be improved by introgression of TP trait in elite wheat cultivars with better source capacity, particularly in winter wheat background. Equivalence for grain weight per spike was not tested; the study may be underpowered to exclude small reductions, and findings are limited to a single glasshouse environment. Multi-environment trials are needed to confirm non-inferiority. In summary, we identified mutations in two genes exhibiting a nearly complete co-segregation with TP phenotype and developed co-dominant functional markers for early generation TP trait selection in hybrid wheat breeding programs (Whitford et al. 2013; Selva et al. 2020; Rohde et al. 2025). We also employed CRISPR-Cas9 gene editing to functionally test a candidate gene representing a direct functional genomics approach to dissect the underlying genetic mechanism. This study advances our understanding on the genetic basis of a nearly half century old scientific riddle and provides candidate functional markers for early-generation TP trait selection in wheat breeding programs.

## Supporting Data

Fig. S1: Procedures of biparental populations development.

Table S1: List of primers used in this study.

## Acknowledgements

We are thankful to Prof. Yuling Jiao for providing lab resources and guidance to conduct the research, Dr. Mingjiu Li for guidance on designing the experiments, and Dr. Yuange Wang for providing WTP wheat material. The infrastructural support received from Peking University Institute of Advanced Agricultural Sciences is also gratefully acknowledged.

## Author Contributions

**QR** conceived the idea of the project and designed the experiments. **ZA** provided germplasm materials. **QR** conducted experiments and performed data analysis. **QS**, **SR** and **ZA** provided inputs in discussion. **QS** provided resources for experiments. **QR** wrote the manuscript with the inputs from the co-authors. All co-authors read and approved the manuscript.

## Data Availability

All data generated or analyzed during this study are included in this manuscript and its supplementary information.

---

**Fig. S1.**
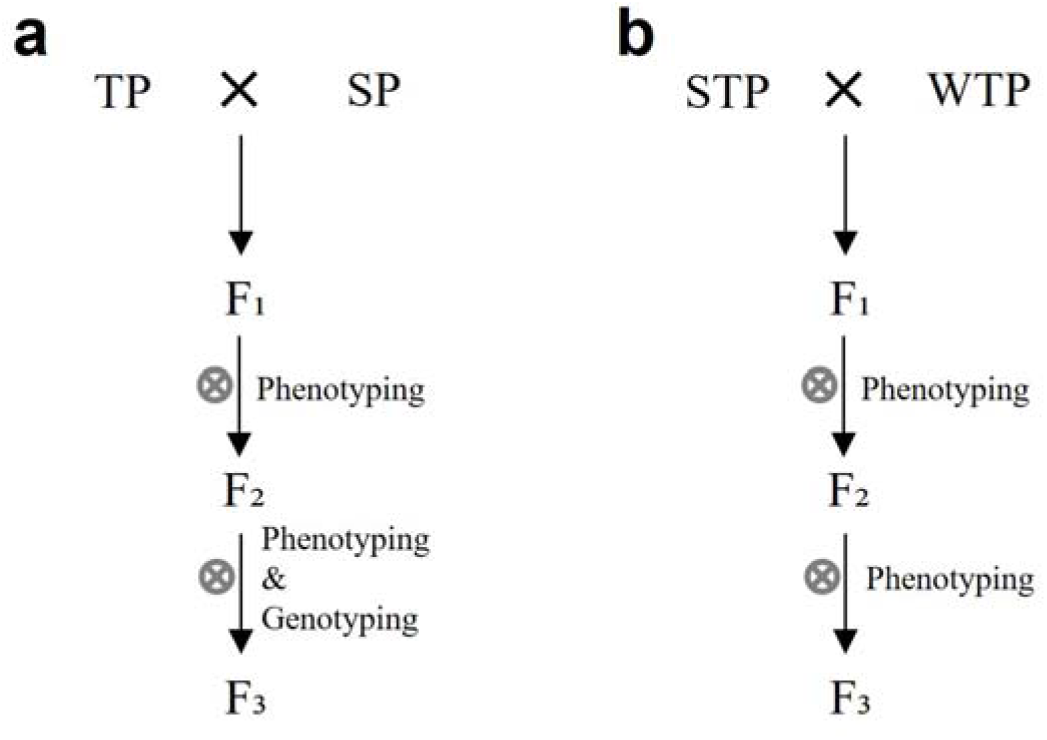
Procedures of biparental populations development. **(a)** Procedure of populations development between triple pistil (TP) and single pistil (SP) wheat accessions. STP wheat was used as female parent, along with Fielder and Pirbak as male parents. (**b**) Procedure of population development between two distinct TP wheat accessions.

**Table S1.**
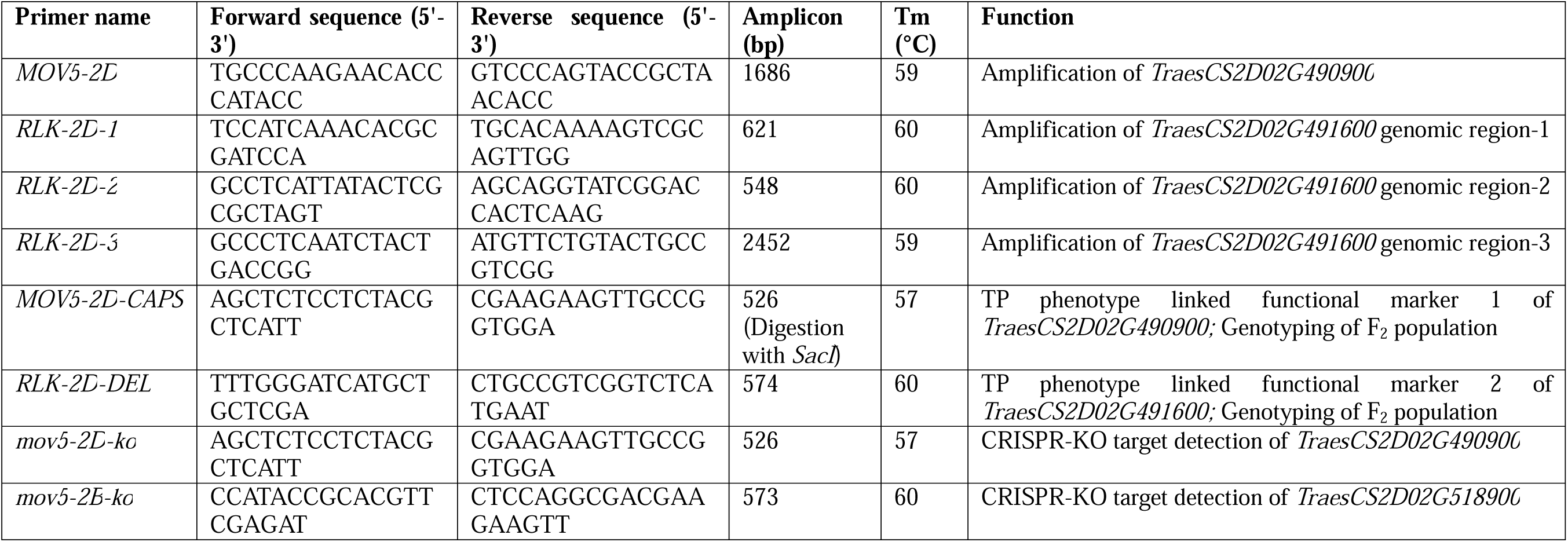
List of primers used in this study. All data generated or analyzed during this study are included in this manuscript and its supplementary information.

## References

Abdulrazzak N, Pollet B, Ehlting J, et al (2006) A coumaroyl-ester-3-hydroxylase insertion mutant reveals the existence of nonredundant meta-hydroxylation pathways and essential roles for phenolic precursors in cell expansion and plant growth. Plant Physiol. 10.1104/pp.105.069690

Adamski NM, Simmonds J, Brinton JF, et al (2021) Ectopic expression of Triticum polonicum VRT-A2 underlies elongated glumes and grains in hexaploid wheat in a dosage-dependent manner. Plant Cell. 10.1093/plcell/koab119

Ali Z, Raza Q, Atif RM, et al (2019) Genetic and molecular control of floral organ identity in cereals. Int J Mol Sci

Andersen JR, Lübberstedt T (2003) Functional markers in plants. Trends Plant Sci

Appels R, Eversole K, Feuillet C, et al (2018) Shifting the limits in wheat research and breeding using a fully annotated reference genome. Science (1979). 10.1126/science.aar7191

Bagge M, Xia X, Lübberstedt T (2007) Functional markers in wheat. Curr Opin Plant Biol

Besseau S, Hoffmann L, Geoffroy P, et al (2007) Flavonoid accumulation in Arabidopsis repressed in lignin synthesis affects auxin transport and plant growth. Plant Cell. 10.1105/tpc.106.044495

Chang TG, Zhu XG (2017) Source-sink interaction: a century old concept under the light of modern molecular systems biology. J Exp Bot

Chen J, Zhang L, Bingli W (1983) A preliminary report on the discovery and breeding of the “trigrain wheat". Acta Agron Sinica 0–10

Cui Y, Lu X, Gou X (2022) Receptor-like protein kinases in plant reproduction: Current understanding and future perspectives. Plant Commun

Debernardi JM, Tricoli DM, Ercoli MF, et al (2020) A GRF–GIF chimeric protein improves the regeneration efficiency of transgenic plants. Nat Biotechnol. 10.1038/s41587-020-0703-0

Erenstein O, Jaleta M, Abdul Mottaleb K, et al (2022) Global Trends in Wheat Production, Consumption and Trade. In: Wheat Improvement: Food Security in a Changing Climate

Hu MJ, Zhang HP, Cao JJ, et al (2016) Characterization of an IAA-glucose hydrolase gene TaTGW6 associated with grain weight in common wheat (Triticum aestivum L.). Molecular Breeding. 10.1007/s11032-016-0449-z

Igrejas G, Branlard G (2020) The importance of wheat. In: Wheat Quality For Improving Processing And Human Health

Ishida Y, Tsunashima M, Hiei Y, Komari T (2015) Wheat (Triticum aestivum l.) Transformation using immature embryos. Methods in Molecular Biology. 10.1007/978-1-4939-1695-5_15

Koppolu R, Chen S, Schnurbusch T (2022) Evolution of inflorescence branch modifications in cereal crops. Curr Opin Plant Biol

Liu S, Lei J, Zhang J, et al (2023) Genome-wide identification and analysis of wheat LRR-RLK family genes following Chinese wheat mosaic virus infection. Front Plant Sci. 10.3389/fpls.2022.1109845

Ma S, Wang M, Wu J, et al (2021) WheatOmics: A platform combining multiple omics data to accelerate functional genomics studies in wheat. Mol Plant

Mahlandt A, Rawat N, Leonard J, et al (2021) High-resolution mapping of the Mov-1 locus in wheat by combining radiation hybrid (RH) and recombination-based mapping approaches. Theoretical and Applied Genetics. 10.1007/s00122-021-03827-w

Peng ZS (2003) A new mutation in wheat producing three pistils in a floret. J Agron Crop Sci. 10.1046/j.1439-037X.2003.00040.x

Peng ZS, Martinek P, Kosuge K, et al (2008) Genetic mapping of a mutant gene producing three pistils per floret in common wheat. J Appl Genet. 10.1007/BF03195606

Raza Q, Ali Z, Karim I, et al (2019) Genetic Analysis of Triple Pistil Wheat Derived F2 Populations to Enhance Genetic Yield Potential. Res Plant Biol. 10.25081/ripb.2019.v9.3752

Rohde A, Albertsen MC, Boden SA, et al (2025) New genomic resources to boost research in reproductive biology to enable cost effective hybrid seed production. Plant Genome 18:e70092. 10.1002/tpg2.70092

Sakuma S, Schnurbusch T (2020) Of floral fortune: tinkering with the grain yield potential of cereal crops. New Phytologist

Schauberger B, Ben-Ari T, Makowski D, et al (2018) Yield trends, variability and stagnation analysis of major crops in France over more than a century. Sci Rep. 10.1038/s41598-018-35351-1

Schoen A, Yoshikawa GV, Sharma PK, et al (2025) WUSCHEL-D1 upregulation enhances grain number by inducing formation of multiovary-producing florets in wheat. Proceedings of the National Academy of Sciences, 122(42), p.e2510889122.

Selva C, Riboni M, Baumann U, et al (2020) Hybrid breeding in wheat: How shaping floral biology can offer new perspectives. Functional Plant Biology

Selva C, Yang X, Shirley NJ, et al (2023) HvSL1 and HvMADS16 promote stamen identity to restrict multiple ovary formation in barley. J Exp Bot. 10.1093/jxb/erad218

Simmonds J, Scott P, Brinton J, et al (2016) A splice acceptor site mutation in TaGW2-A1 increases thousand grain weight in tetraploid and hexaploid wheat through wider and longer grains. Theoretical and Applied Genetics. 10.1007/s00122-016-2686-2

Skovmand B, Reynolds MP, Delacy IH (2001) Searching genetic resources for physiological traits with potential for increasing yield. In: Reynolds MP, Ortiz-Monasterio JI, McNab A (eds) Application of physiology in wheat breeding. CIMMYT, Mexico, pp 124–135

Sreenivasulu N, Schnurbusch T (2012) A genetic playground for enhancing grain number in cereals. Trends Plant Sci

Tadesse W, Sanchez-Garcia M, Assefa SG, et al (2019) Genetic Gains in Wheat Breeding and Its Role in Feeding the World. Crop Breed Genet Genom. 10.20900/cbgg20190005

Wang Y, Du F, Wang J, et al (2022) Improving bread wheat yield through modulating an unselected AP2/ERF gene. Nat Plants. 10.1038/s41477-022-01197-9

Wang Z, Xu D, Ji J, et al (2009) Genetic analysis and molecular markers associated with multi-gynoecia (Mg) gene in Trigrain wheat. Canadian Journal of Plant Science. 10.4141/cjps09022

Whitford R, Fleury D, Reif JC, et al (2013) Hybrid breeding in wheat: Technologies to improve hybrid wheat seed production. J Exp Bot. 10.1093/jxb/ert333

Yamamoto N, Chen Z, Guo Y, et al (2023) Gene co-expression modules behind the three-pistil formation in wheat. Funct Integr Genomics 23:. 10.1007/s10142-023-01052-w

Yang Z, Chen Z, Peng Z, et al (2017) Development of a high-density linkage map and mapping of the three-pistil gene (Pis1) in wheat using GBS markers. BMC Genomics. 10.1186/s12864-017-3960-7

Yu M, Nie X, Ke S, et al (2025) Receptor-like kinase cleavage: molecular mechanism and regulatory functions in plants. New Phytologist 246:2478–2483. 10.1111/nph.70174

Yu ZY, Luo Q, Peng Z, et al (2021) Genetic mapping of the three-pistil gene Pis1 in an F2 population derived from a synthetic hexaploid wheat using multiple molecular marker systems. Cereal Res Commun 49:31–36. 10.1007/s42976-020-00078-1

Zhang D, Tang S, Chen J, et al (2025) Chromosomal inversion at the DG1 promoter drives double-grain spikelets and enhances grain yield in sorghum. Nat Plants 453–467. 10.1038/s41477-025-01937-7

Zhang X, Jia H, Li T, et al (2022) TaCol-B5 modifies spike architecture and enhances grain yield in wheat. Science (1979). 10.1126/science.abm0717

Zhang X, Meng W, Liu D, et al (2024) Enhancing rice panicle branching and grain yield through tissue-specific brassinosteroid inhibition. Science (1979). 10.1126/science.adk8838

Zhou J, Li W, Yang Y, et al (2023) A promising QTL QSns.sau-MC-3D.1 likely superior to WAPO1 for the number of spikelets per spike of wheat shows no adverse effects on yield-related traits. Theoretical and Applied Genetics. 10.1007/s00122-023-04429-4

Zhu X, NI Y jing, HE R shi, et al (2019) Genetic mapping and expressivity of a wheat multi-pistil gene in mutant 12TP. J Integr Agric. 10.1016/S2095-3119(18)61935-5

